# Spore-Based Biocomposite Thermoplastic Polyesters with Enhanced Toughness and Programmable Disintegration

**DOI:** 10.64898/2026.02.24.707801

**Authors:** Han Sol Kim, Emily Fan, Arthi Chandra, Evangeline Meyler, Joanne Tang, Myung Hyun Noh, Adam M. Feist, Jonathan K. Pokorski

## Abstract

Thermoplastic polyesters are widely used in commodity and high-performance applications due to their tunable and exceptional properties, versatile performance, and increasing relevance in sustainable materials. Integrating biological functionality into these polymers offers a promising route to enhance performance and end-of-life behavior beyond what conventional additives can achieve. Here, we report the generalization of an embedded spore-based engineered living material concept to three representative thermoplastic polyesters; polycaprolactone (PCL), polylactic acid (PLA), and poly(butylene adipate-co-terephthalate) (PBAT). Heat-shock-tolerized *Bacillus subtilis* spores were compounded with each polyester as a living biofiller via hot melt extrusion. The resulting biocomposite polyesters retained high spore viability (>90%) after extrusion and exhibited improved mechanical performance (up to 41% toughness improvement compared to neat polymers). End-of-life behavior was evaluated in a microbially-limited composting environment, where spore-containing PCL exhibited nearly complete disintegration within five months, corresponding to a ∼7-fold increase in degradation kinetics relative to neat PCL. Finally, 3D printing of biocomposite PCL was demonstrated through fused deposition modeling and direct ink writing methodologies. Together, this work demonstrated the successful extension of spore-based engineered living materials from thermoplastic polyurethane to multiple thermoplastic polyesters.

**Graphical Abstract:** 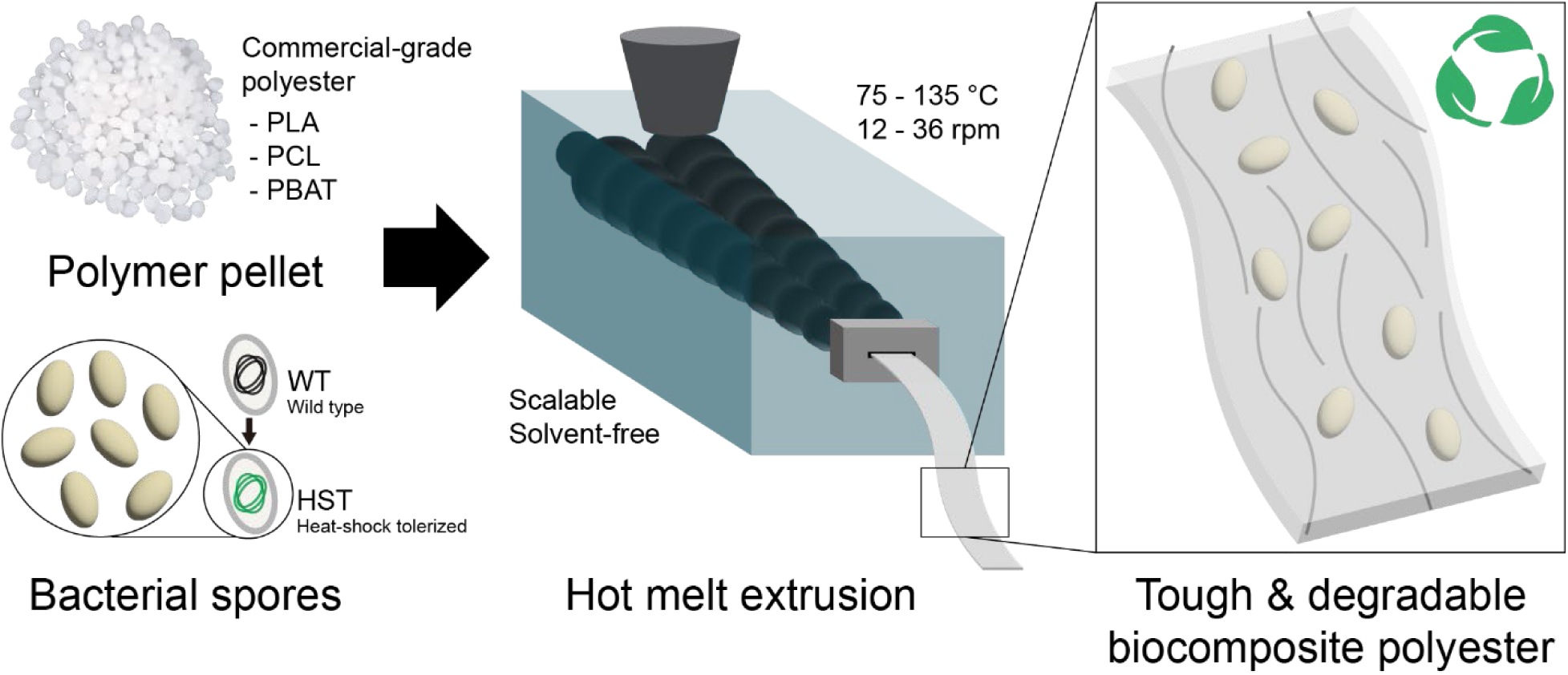

## 1. Introduction

Polyesters are one of the most important classes of polymers with extensive applicability across diverse industrial sectors.^1,2^ Polyesters contain repeating ester bonds in their polymeric main chains and can range from soft and extensible to stiff and rigid based on monomer composition.^3^ Specifically, thermoplastic polyesters soften and can be processed upon heating and solidify by cooling without undergoing significant chemical change, offering exceptional manufacturing versatility.^4^ This manuscript focuses on three extensively used polyesters: polylactic acid (PLA), polycaprolactone (PCL) and poly(butylene adipate-co-terephthalate) (PBAT). PLA and PCL are used across various sectors including additive manufacturing and biomedical engineering.^5–8^ Whereas, PBAT features high flexibility and toughness and is used in packaging and agricultural industries.^9,10^ Overall, thermoplastic polyesters occupy a central role in both commodity and high-performance applications, bridging the gap between conventional plastics and advanced engineering materials.

To advance the functionalities of traditional polymers, there have been efforts to incorporate novel additives into the polymer matrix.^11–13^ Currently, some of the most innovative additives to generate novel polymer properties are living cells as they possess inherent abilities for both self-regeneration and self-replication, while responding to external environmental stimuli. So called engineered living materials (ELMs) pair living cells with polymers to develop biofunctional materials that can outperform conventional polymers.^14–16^ Our previous research has shown that bacterial spores embedded within a thermoplastic polyurethane (TPU) via melt extrusion improved the tensile properties of the composite material as well-defined and well-dispersed submicron reinforcing fillers inside the TPU matrix, and facilitated disintegration of plastic at its end of life.^17^ In addition, bacterial spores prevented thermal oxidation of TPU, significantly improving its reprocessability.^18^

Building on our previous work, we envisioned that the concept of a spore-bearing biocomposite polymer can be expanded to other polymers beyond TPUs. Herein, we explore the generalization of spore-based biocomposite polymers using three common thermoplastic polyesters, PLA, PCL and PBAT (**Figure 1**). Mechanical and biological compatibility between spores and the three polyesters were investigated based on the tensile properties of the composite materials and spore survivability post compounding, respectively. Disintegration of biocomposite polyesters was monitored in microbially-depleted compost. Finally, a potential application of a biocomposite polymer was demonstrated through 3D printing using filaments of spore-containing polyester.

**Figure 1.**
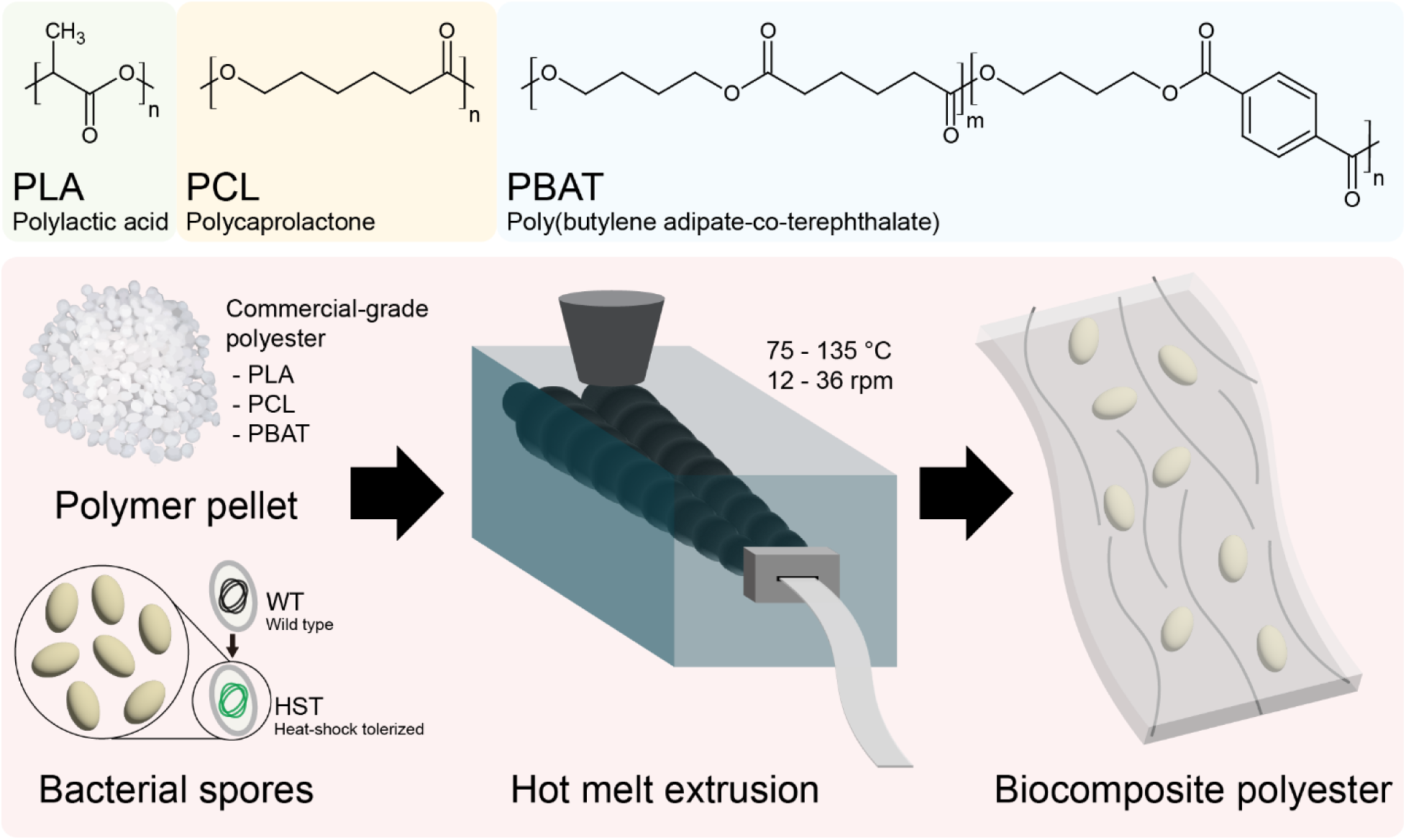
Schematic illustration of the fabrication of biocomposite polyesters filled with bacterial spores. Heat shock tolerized (HST) B. subtilis spores (ATCC 6633 HST strain) are used as biofillers for PLA, PCL and PBAT polyesters.

## 2. Results & Discussion

### 2.1. Fabrication and Characterization of Biocomposite PCL, PLA and PBAT

We reasoned that the use of an evolutionarily engineered *B. subtilis* strain (ATCC6633 HST), developed in our previous work, could effectively be employed as a biological component of multiple additional biocomposite polyesters.^17^ This was motivated by our preliminary studies, where the ATCC 6633 HST exhibited strong compatibility with a polyester-based TPU and demonstrated pronounced synergy as a biocomposite material. Specifically, ATCC 6633 HST showed a favorable TPU assimilation rate and high resistance to elevated temperature and shear, while remaining viable during the melt processing of polymer. Thus, we moved forward with this evolved strain as a filler for the polyesters PCL, PLA and PBAT.

To evaluate the thermal tolerance of the ATCC 6633 HST strain, spores were compounded with a model polymer (TPU) at different processing temperatures to determine the maximum allowable heat tolerance to maintain spore viability (**Figure S1**). The spores retained approximately 90% viability up to 140 °C, after which heat-induced inactivation became apparent. Following processing at 145 °C, spore viability after incorporation decreased to approximately 49% of the control and exhibited a logarithmic decline with further increases in processing temperature. Based on these results, commercial polyesters that could be extruded below 140 °C were selected for analysis (PCL: Sigma-Aldrich 704105, PLA: NatureWorks Ingeo^TM^ 4950D, PBAT: BASF Ecoflex® F Blend C1200). Optimized processing temperatures of the selected PCL, PLA and PBAT materials in a twin screw, corotating microcompounder were 65 °C, 120 °C and 135 °C, respectively, all of which are below the temperature threshold for high ATCC 6633 HST spore viability. Processing parameters, such as mixing speed, mixing time and extrusion speed, were further optimized for each polyester (**Table S1**) to yield uniform and smooth biocomposite polyester ribbons (**Figure 2**). Spore loading for each biocomposite polyester was 0.5% (w/w) based on optimal loading in TPU for improving mechanical properties,^18^ and visual inspection resulted in the conclusion that incorporation of spores did not significantly affect the appearance of the extrudates or the processability of the polyesters (**Figure 2**).

**Figure 2.**
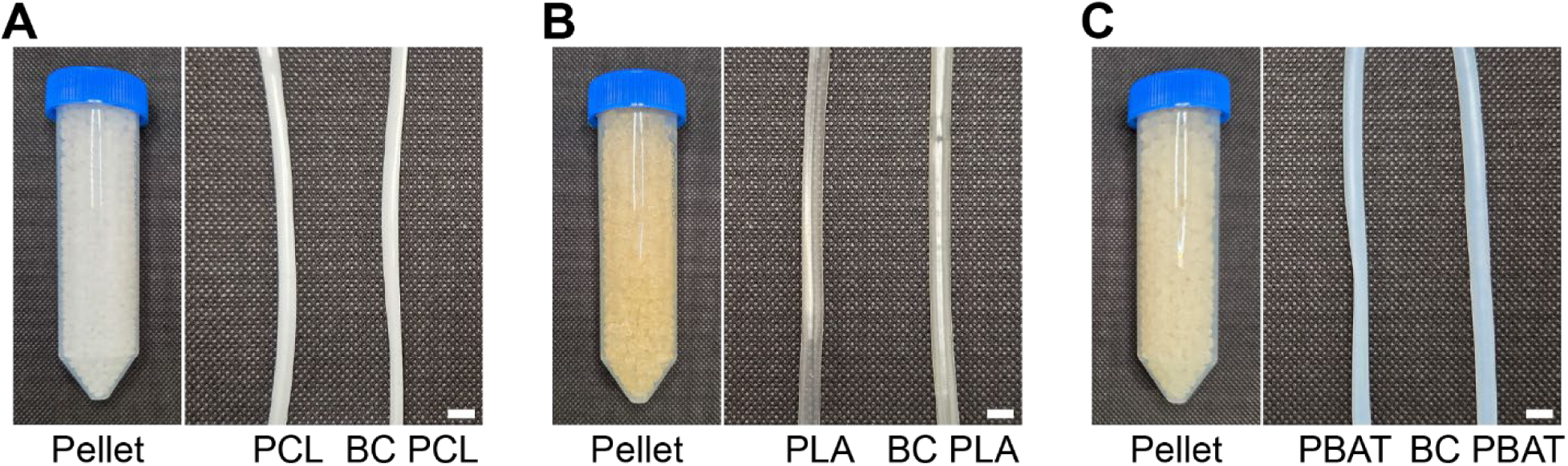
Photographs of polyesters with (‘BC’) and without spores. Pellets, in tubes, left and processed ribbon shape extrudates, right were processed using a lab-scale extruder (scale bars = 1 cm). (A) PCL, (B) PLA and (C) PBAT polyesters.

The viability of spores embedded in the biocomposite polyesters was quantified by extracting the spores from the composites by dissolving the polymer component of the composites using an appropriate organic solvent (tetrahydrofuran for PBAT and *N,N*-dimethylformamide for PCL and PLA). Spores were further washed with water and collected via centrifugation. Additionally, Neat spores were subjected to the same solvent treatment protocol and their viabilities were used as baselines. Spores retained, on average, 96%, 90% and 95% viability in biocomposite PCL, PLA and PBAT, respectively (**Figure 3**). Overall, this analysis demonstrated that the spores showed nearly full survivability after compounding with polyesters, regardless of polymer type. These results also indicated that the spores were compatible with all three polyesters tested, in the context that the processing temperature remained within below the thermal tolerance threshold of high spore viability (< 145 °C) (**Figure S1**).

**Figure 3.**
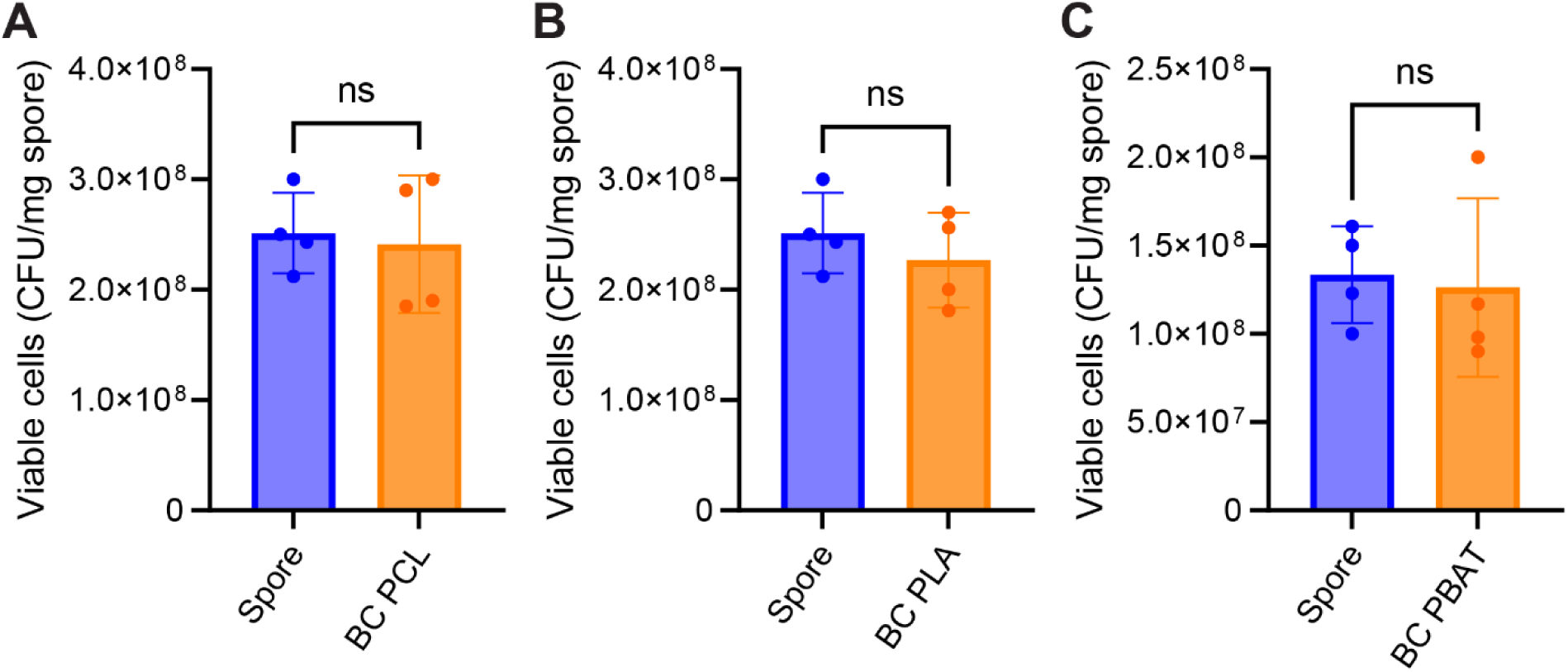
Viability of spores before and after compounding with PCL (A), PLA (B) and PBAT (C). Baseline spores were subjected to the same solvent extraction processes for spores in biocomposite polyesters. Data are presented as mean values ± standard deviations from four independent experiments. Welch’s t test was performed for statistical analysis. P < 0.05 was considered statistically significant. ns: not significant.

After confirming spore viability in biocomposite polyesters, the physical properties of the materials were investigated. For tensile testing, biocomposite polyester ribbons were reshaped into dogbone geometries (ISO 527-2-5B) using a die cutting method. Tensile properties of the were evaluated by subjecting the specimens to uniaxial tensile loading until fracture. The resulting stress–strain data were used to determine four key tensile properties such as toughness, ultimate tensile strength, elongation at break, and Young’s modulus (**Figures 4 and S2-3**). The incorporation of spores enhanced the toughness of all polyesters tested. Specifically, the toughness of PCL, PLA, and PBAT increased by 33%, 42%, and 31%, respectively, upon spore addition, with statistical significance. These results are consistent with our previous findings for TPU-based composites and suggest effective interfacial adhesion between the spores and the polymer chains.^17,19^ Water contact angle analyses of biocomposite polyesters showed a significant increase in hydrophilicity of composites compared to neat polymers without spores. Even though a small amount of spores (0.5 wt%) was added to polyesters, polyesters are highly crystalline and inherently hydrophobic (**Figure S4**). In this regard, the improved tensile properties of polyesters by spore incorporation can be attributed to the hydrogen bonding and/or dipole-dipole interaction between surface functional groups of spores and the ester bonds in polyesters.^20^ DSC and FTIR exhibited no significant thermal or chemical change between the neat polyesters and biocomposite polyesters (**Figures S5-S6**)

**Figure 4.**
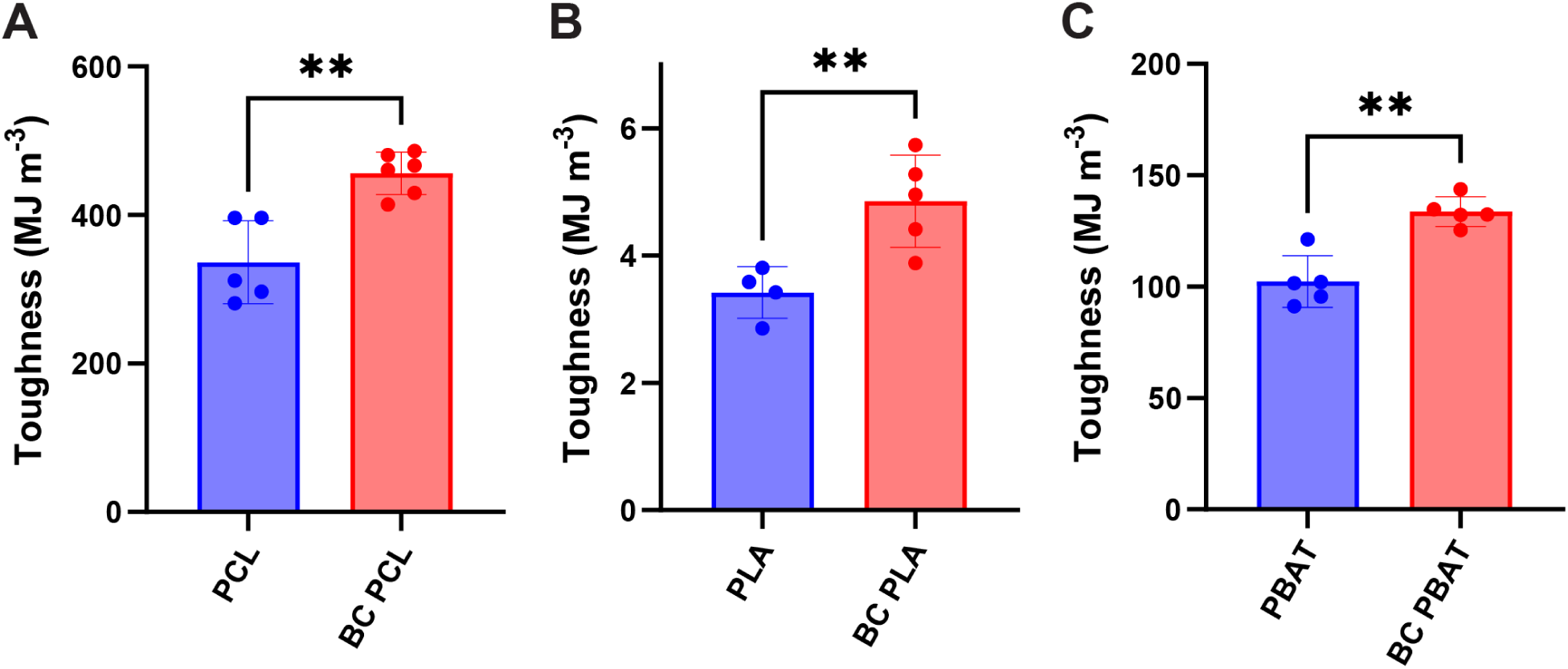
Toughness of polyesters with and without spores. Data are presented as mean values ± standard deviations from four or more independent experiments. Welch’s t test was performed for statistical analysis. P < 0.05 was considered statistically significant. Detailed indicators are as follows – ns: not significant; *P < 0.05; **P < 0.01; ***P < 0.001; ****P < 0.0001.

### 2.2. Disintegration of Biocomposite Polyesters

End-of-life disintegration tests were performed to characterize the breakdown of the formulated biocomposites. To do so within a limited timeframe, neat and biocomposite polyesters were hot-pressed to increase their surface area and, thereby, potentially accelerate disintegration. Ribbon shaped samples (200 mg) were pressed between two parallel plates under 6.3 MPa of pressure at elevated temperatures (PCL: 55 °C / PLA: 100 °C / PBAT: 80 °C). The hot-pressed polyesters with and without incorporated bacterial spores were incubated in autoclaved compost to assess disintegration of polymer samples under simulated microbially limited end-of-life conditions (**Figure 5A**). It was found that previously, autoclaved compost suppressed background microbial activity enabling isolation of degradation driven by spore embedment and likely ATCC6633 HST-driven hydrolysis.^17^ Incubation was conducted at 37 °C and 45–55% relative humidity (**Figure 5B**).

**Figure 5.**
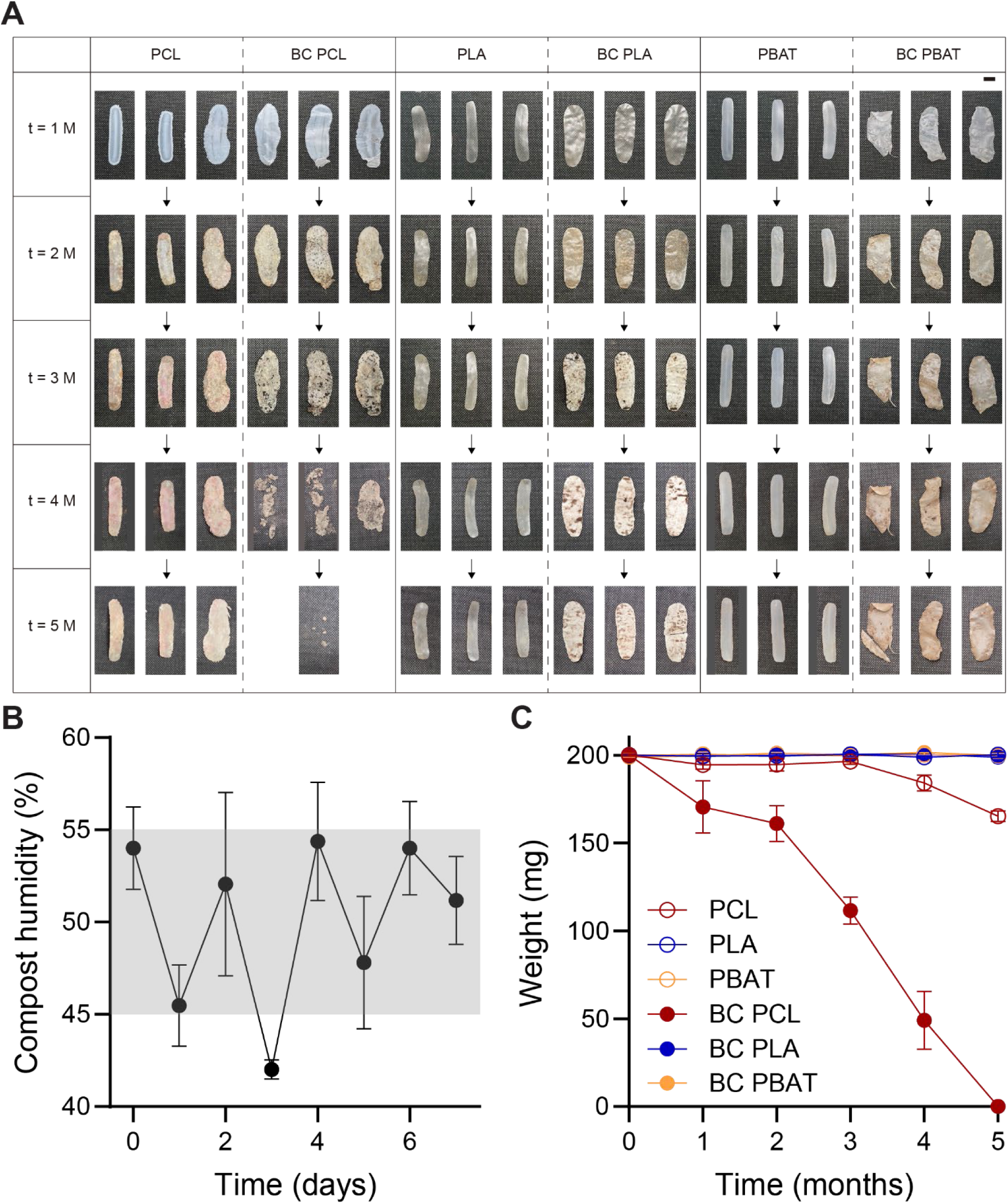
Disintegration of polyesters with (‘BC’) and without spores in autoclaved compost during 5 months of incubation at 37 °C a 45–55% relative humidity. (A) Photographs of polyester samples. (B) Compost humidity was maintained at 45-55 % after a few days of conditioning. (C) Weight change of the examined polyesters samples during the compost incubation.

Disintegration tests revealed polymer-specific divergent extents of degradation. PBAT and PLA showed no visible disintegration or measurable mass loss over 5 months, regardless of spore incorporation, retaining ∼100% of their initial mass. In contrast, neat PCL exhibited ∼17% mass loss after 5 months, while spore-containing biocomposite PCL underwent near-complete disintegration (∼100% mass loss) over the same period. Although PBAT and PLA are classified as biodegradable/compostable,^21,22^ autoclaved compost provided no apparent microbial and enzymatic activity for their degradation, and hydrolytic chain scission under these conditions was minimal. Spore-containing PBAT and PLA samples exhibited more significant surface-level deterioration, including transparency loss, discoloration, and cracking, compared to neat polyesters, but it did not necessarily lead to corresponding mass loss. Such changes on biocomposite PBAT and PLA can be possibly explained by the colonization of germinated ATCC6633 HST spores or other recolonizing microbes in/on plastic surfaces.^23^ However, ATCC6633 HST was not specifically selected or evolutionarily engineered to degrade PBAT or PLA and could not effectively facilitate its degradation likely due to the absence of suitable enzymes or limited accessibility of bacterial enzymes to these specific ester bonds.^24^ In contrast, the ATCC 6633 HST strain apparently functioned as an efficient PCL degrader, exhibiting approximately 700% faster degradation kinetics, calculated from the slope of the mass-loss curves, for biocomposite PCL relative to neat PCL. Mass loss of the neat PCL over the incubation test can be explained by hydrolysis or repopulation of the compost environment by additional active microbes and subsequent degradation. This finding suggests more favorable ester bond accessibility in PCL compared to PLA or PBAT despite its relatively low ester bond concentration.^3^

### 2.3. 3D Printing of Biocomposite PCL

PCL is regarded as a biocompatible, non-toxic and cost-effective polymer^25^. 3D printing is one of the most prominent applications of PCL. Both fused deposition modeling (FDM) and direct ink write (DIW) printing were demonstrated using biocomposite PCL. For FDM, biocomposite PCL was fabricated into filament form by simply replacing the extruder exit die with a circular die (2.0 mm diameter). For DIW printing, biocomposite PCL pieces were loaded into an aluminum syringe and extruded under pneumatic pressure at an elevated temperature. Then, the molten biocomposite PCL was dispensed through a nozzle (0.5 mm diameter) using pneumatic pressure and deposited onto a print bed. Biocomposite PCL showed printability in both FDM and DIW printing modalities, indicating that spore incorporation does not interfere with this major application of PCL (**Figure 6**). It should be noted that *B. subtilis* is non-pathogenic and non-toxic bacterium, and is generally recognized as safe (GRAS) by the FDA.^26^

**Figure 6.**
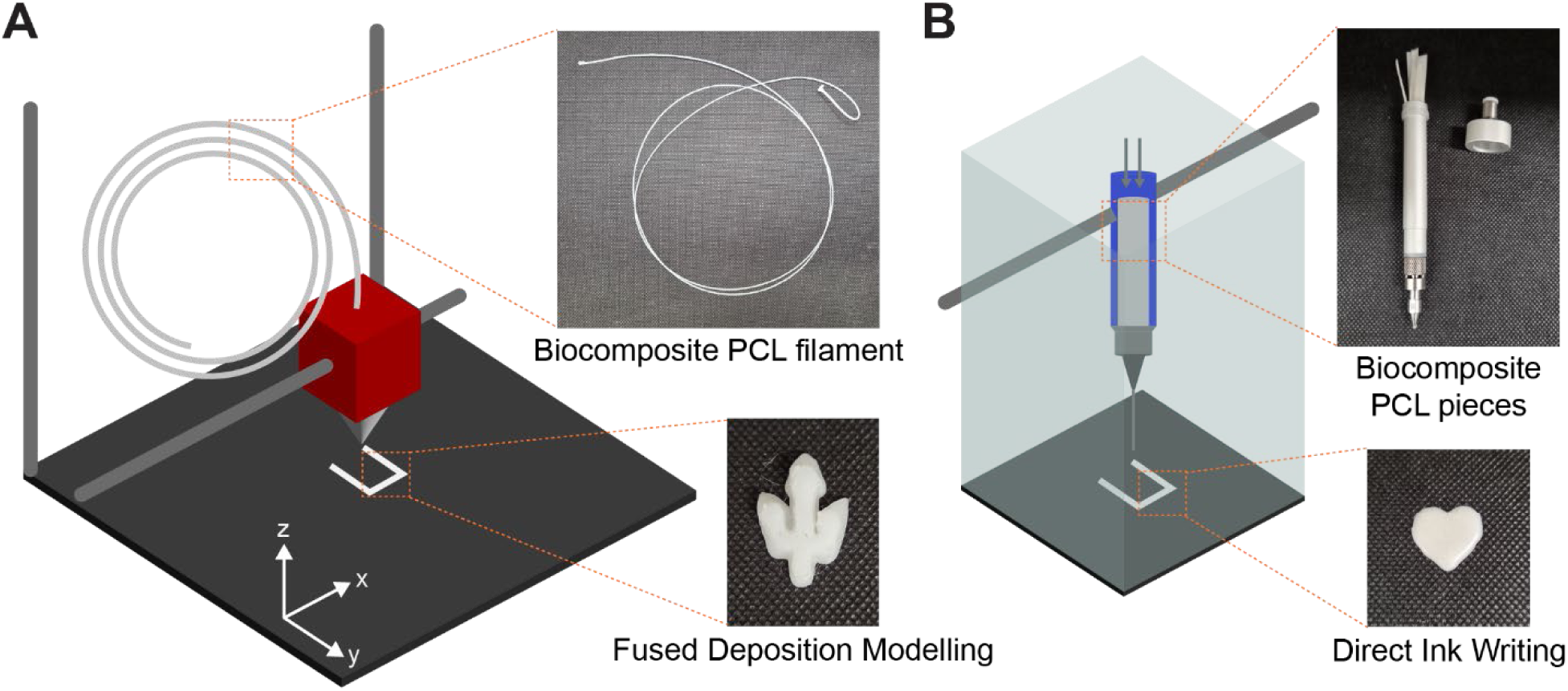
Demonstration of 3D printing using biocomposite PCL via fused deposition modeling (A) and direct ink writing (B).

Viability of the embedded spores in biocomposite PCL after 3D printing was quantified (**Figure S7**). Spores retained 45% and 83% viability after FDM- and DIW-based 3D printing, respectively. Even though FDM was carried out at a very high nozzle temperature (200 ℃), the relatively short contact between the biocomposite PCL filament and heated nozzle resulted in a reasonable retainment of spore viability after printing. Spores maintained high viability after DIW printing, likely, due to its mild printing condition (130 ℃, 400 kPa).

## 3. Conclusions

In this work, we demonstrated the successful extension of spore-based engineered living materials from TPUs to multiple thermoplastic polyesters, including PCL, PLA, and PBAT. By matching the polymer processing conditions with the thermal tolerance thresholds of heat shock tolerized *B. subtilis* spores, biocomposite polyesters were fabricated via conventional melt extrusion while retaining high spore viability. Across all three polyester systems, the incorporation of a small fraction of spores (0.5% (w/w)) enhanced tensile toughness and increased surface hydrophilicity, indicating favorable mechanical and interfacial compatibility between spores and polyester matrices. The high viability of the embedded spores shows great promise in having spores that can germinate and be programmed to carry out biological processes to not only enhance degradation but enable new functionalities.

End-of-life studies under microbially-limited composting conditions revealed polymer-specific disintegration behavior. PLA and PBAT and their biocomposite formulations showed limited mass loss. Notably, biocomposite PCL exhibited nearly complete disintegration within five months, with degradation kinetics approximately seven times faster than neat PCL. This result underscores the ability of embedded spores to actively drive polymer disintegration in otherwise unfavorable environments.

Importantly, the biocomposite polyesters maintained excellent processability, enabling both filament-based fused deposition modeling and extrusion-based 3D printing of spore-containing PCL. These results demonstrate that bacterial spores can serve not only as a versatile reinforcing agent for multiple polyesters, but also as a biofunctional degrader for some polyesters. This platform provides a versatile and scalable strategy for designing next-generation living plastics with enhanced mechanical performance, reprocessability, and programmable end-of-life behavior, and can be further expanded through strain engineering tailored to specific polymer chemistries and applications.

## 4. Methods

### 4.1. Cells, materials and reagents

#### Cells

Evolutionarily engineered *B. subtilis* (A5_F40_I1 strain, ATCC 6633 HST) developed in our previous work was used as a biofiller of polyesters.^17^

#### Materials

Soft-grade, polyester-based thermoplastic polyurethane pellets (TPU, Elastollan® BCF45) and polybutylene adipate terephthalate (PBAT, Ecoflex® F Blend C1200) were gifted from BASF (Wyandotte, MI, USA). Polycaprolactone (PCL, M_n_ 45 kDa) was purchased from Sigma-Aldrich (St. Louis, MO, USA). Polylactic acid (PLA, Ingeo^TM^ 4950D) was purchased from NatureWorks (Plymouth, MN, USA). Compost for disintegration test was gifted from Prof. Jason Locklin and Dr. Evan White (New Materials Institute, University of Georgia, Athens, GA, 30602, USA)

#### Reagents

Luria–Bertani (LB) powder (Miller), BD Difco^TM^ nutrient broth, BD Bactor^TM^ agar and 10x phosphate-buffered saline (PBS) pH 7.4 were purchased from Fisher Scientific (Waltham, MA, USA). KCl, MgSO_4_, CaCl_2_, FeSO_4_, MnCl_2_, lysozyme, tetrahydrofuran (THF) and *N,N*-dimethylformamide (DMF) were purchased from Sigma–Aldrich.

### 4.2. Spore production

Spores were produced by following our previously reported protocols.^18^ LB medium was prepared by dissolving LB powder in deionized (DI) water at 25 g L^-1^ concentration. Difco sporulation medium (DSM) was prepared by dissolving BD Difco^TM^ nutrient broth (8 g L^-1^), KCl (1 g L^-1^) and MgSO_4_ (0.12 g L^-1^) in DI water. Both media were autoclaved at 121 °C for 20 min. For DSM, FeSO_4_ (1 nM) CaCl_2_ (1 mM) and MnCl_2_ (1 µM) were added after autoclaving.

ATCC 6633 HST glycerol stock frozen at-80 °C was thawed at room temperature, followed by inoculation in LB medium at 1 % (v/v). After incubation at 37 °C with 250 rpm of shaking overnight, the seed culture was inoculated into DSM at 1 % (v/v) for the main culture and subsequent sporulation. DSM culture was incubated at 37 °C at 250 rpm shaking for an extended period of the time (2 days) to induce sporulation. Crude spore suspension was centrifuged at 12,000 g at 4 °C for 20 min (Avanti J-E, Beckman Coulter Life Science, Brea, CA, USA). Spores were purified via repetitive washing using fresh PBS (100 mM, pH 7.4, sterile); (i) resuspension in PBS, (ii) centrifugation at 2200 g at 25 °C for 10 min (5810 R, Eppendorf) and (iii) supernatant removal. After three washes, remaining vegetative cells in the suspension were lysed using lysozyme enzyme (2.5 mg mL^-1^) at 37 °C at 250 rpm shaking for 1 h. Three washes were followed by lysozyme treatment. Spores were further heat-treated at 65 °C under non shaking condition for 1 h. After three washes using DI water, spore suspensions were frozen using liquid nitrogen and then lyophilized for 3 days (FreeZone 2.5, Labconco, Kansas City, MO, USA)

### 4.3. Hot melt extrusion

Polymers were processed using a HAAKE^TM^ Mini CTW (Thermo Fisher Scientific), a lab scale conical twin screw extruder, equipped with either a slit exit die (slit size: 5.0 mm x 0.7 mm) or circular die (diameter: 2 mm). ∼5 g of polymer pellets were introduced to the extruder and cycled through the heated recirculation chamber (PCL: 65 °C, 36 rpm / PLA: 135 °C, 36 rpm / PBAT: 120 °C, 36 rpm) for 5 min to melt polyesters. 0.5 % (w/w) of lyophilized spore powder was introduced to the extruder, followed by an additional 15 min of compounding. Screw speed was adjusted to the extrusion speed of each polyester (PCL: 12 rpm / PLA: 18 rpm / PBAT: 24 rpm), and the polyesters were extruded through the exit die. Polyesters without spores were prepared by following the same protocol but without the addition of spores.

### 4.4. Spore viability test

To quantify spore viability inside the biocomposite polyester, 200 mg of biocomposite polyester was diced into small cubes (∼1 x 1 x 1 mm^3^). Biocomposite polyester pieces were soaked in specific solvents (PCL & PLA: DMF / PBAT: TFH) to dissolve the polymer component of the composite materials. Samples were incubated at 40 °C under 200 rpm magnet stirring condition for 30 min. Spores were pelletized by centrifugation at 2200 g at 25 °C for 10 min (5810 R, Eppendorf). Supernatant was removed and fresh solvents were added to each sample (PCL & PLA: DMF / PBAT: TFH), followed by 10 min of rocking at 50 rpm at room temperature. After the final centrifugation, spores were suspended in 100 mM PBS pH 7.4.

Spore suspensions were serially diluted in PBS and viability was determined using a colony forming unit (CFU) assay. For the CFU assay, LB agar plates were prepared. LB powder (25 g L^-1^) and agar (15 g L^-1^) were mixed in DI water and the mixture was autoclaved at 121 °C for 20 min. The solution was aliquoted into petri dishes and cooled at room temperature until gelation. Spore suspension extracted from biocomposite polyesters was applied to the surface of LB agar plates, spread evenly across the surface and incubated at 30 °C for overnight. Number of colonies appeared on the surface of LB agar plates were manually counted.

To determine the viability of baseline spores, pristine spores were subjected to the same solvent extraction process with the spores extracted from biocomposite polyesters, and viability was quantified using CFU assay. Spore survivability was calculated by the ratio of the viability of spores extracted from the biocomposite polyester and baseline spores.

### 4.5. Tensile testing

Ribbon shaped polyester extrudates were tailored into dogbone shapes (ISO 527-2-5B) using a die cutting method. Hot press with 6.3 MPa hydraulic pressure was used for the die cutting. Polyesters were pressed at elevated temperatures (PCL: 55 °C / PLA: 100 °C / PBAT: 80 °C) for 10 min to prevent fracturing during die cutting.

Uniaxial stress was applied to the dogbone specimen using a universal testing machine (Instron 5982, Norwood, MA, USA) equipped with a 100 N load cell at 50 mm/min displacement rate. Stress and strain applied to the specimen was recorded and analyzed to calculate tensile properties, such as toughness, ultimate tensile stress, elongation at break and Young’s modulus.

### 4.6. Water contact angle analysis

Water drops were applied onto the surface of hot-pressed polyesters with and without spores and the contact angle at the surface was measured using contact angle goniometer (Rame-Hart 500, Succasunna, NJ, USA).

### 4.7. Differential scanning calorimetry (DSC)

Discovery DSC 2500 (TA Instruments, New Castle, DE, USA) was used for DSC analysis. Temperature was cycled between-90 and 250 °C at 10 °C min^-1^ rate.

### 4.8. Attenuated total reflectance-Fourier Transform Infrared Spectroscopy (ATR-FTIR)

ATR-FTIR spectra were collected using Nicolet™ iS50 spectrometer (Thermo Fisher Scientific) over the wavenumber range of 4000–525 cm⁻¹ with a spectral resolution of 4 cm⁻¹ at room temperature. Each spectrum was obtained by averaging 16 co-added scans. A potassium bromide (KBr) beam splitter and a deuterated triglycine sulfate (DTGS) detector were employed.

### 4.9. Disintegration test

Hot-pressed polyester specimens (200 mg) with and without spores were subjected to a mass loss–based disintegration test. Hot pressing was carried out by using the same equipment employed for the preparation of dogbone specimens, but without a die cut. Instead, samples were wrapped in aluminum foil. PCL, PLA and PBAT samples were pressed at 55 °C, 100 °C and 80 °C, respectively, for 10 min under 6.3 MPa pressure.

For each sample group, three specimens were prepared and incubated in compost at 37 °C and 45–55% relative humidity for 5 months. Relative humidity was regularly monitored by aliquoting approximately 1 g of compost and measuring its weight before and after overnight drying at 100 °C under vacuum. Specimens were retrieved from the compost monthly for weight measurements and photographic documentation. Prior to imaging, specimen surfaces were gently wiped with autoclaved wipes. The specimens were then dried overnight under ambient conditions before weighing and subsequently returned to the compost for further incubation.

### 4.10. Fused deposition model (FDM) printing

Biocomposite PCL fabricated into filaments with a diameter of 2 mm was used for 3D printing (Prusa, Newark, DE, USA). The nozzle and bed temperatures were set to 200 °C and 25 °C, respectively. The maximum volumetric speed (11 mm³ s⁻²), infill density (15%), and filament diameter (1.6 mm) were adjusted in the slicer software to improve printing resolution.

### 4.11. Direct ink write (DIW) printing

DIW printing of biocomposite PCL was carried out using an Inkredible+ Bioprinter (Cellink, Palo Alto, CA, USA). An aluminum syringe filled with biocomposite PCL pieces was heated to 130 °C, while the printing bed was maintained at 55 °C. The molten biocomposite PCL was deposited at a pressure of 400 kPa.

### 4.12. Statistical analysis

GraphPad Prism 10.6.1 was used for statical analysis. Welch’s t test was performed. *P* < 0.05 was considered statistically significant. Detailed indicators are as follows – ns: not significant; \**P* < 0.05; \*\**P* < 0.01; \*\*\**P* < 0.001; \*\*\*\**P* < 0.0001.

## Author Contributions

The manuscript was written through contributions of all authors. All authors have given approval to the final version of the manuscript.

## Acknowledgements

This work was supported by UC San Diego Materials Research Science and Engineering Center (UCSD MRSEC) (# DMR-2011924) and by the National Cancer Institute (NCI), National Institutes of Health (NIH), Department of Health and Human Services (# 1R01CA293936-01). This work was also sponsored by funding from BOTTLE^TM^ consortium supported by the U.S. Department of Energy’s (DOE’s) Office of Energy Efficiency and Renewable Energy (EERE) and Advanced Manufacturing Office (AMO) (# DE-EE0009296). The authors thank Prof. Jason Locklin and Dr. Evan White (New Materials Institute, University of Georgia, Athens, GA, 30602, USA) for providing PLA and compost.

