## Supplementary Information for "Spore-Based Biocomposite Thermoplastic Polyesters with Enhanced Toughness and Programmable Disintegration"

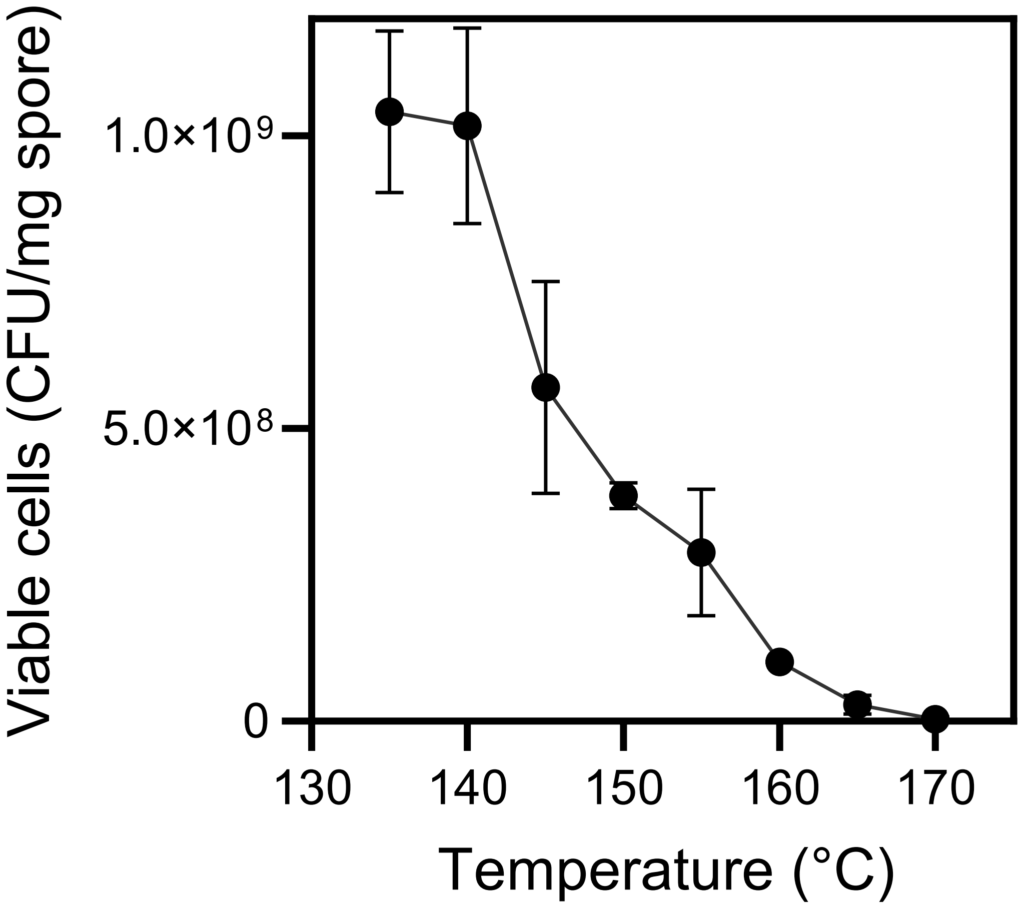

***Figure S1.*** *Viability of spores after being embedded in a biocomposite polymer compounded at different temperatures using a lab scale extruder. Thermoplastic polyurethane was used as a model polymer due to its wide processing temperature range.*

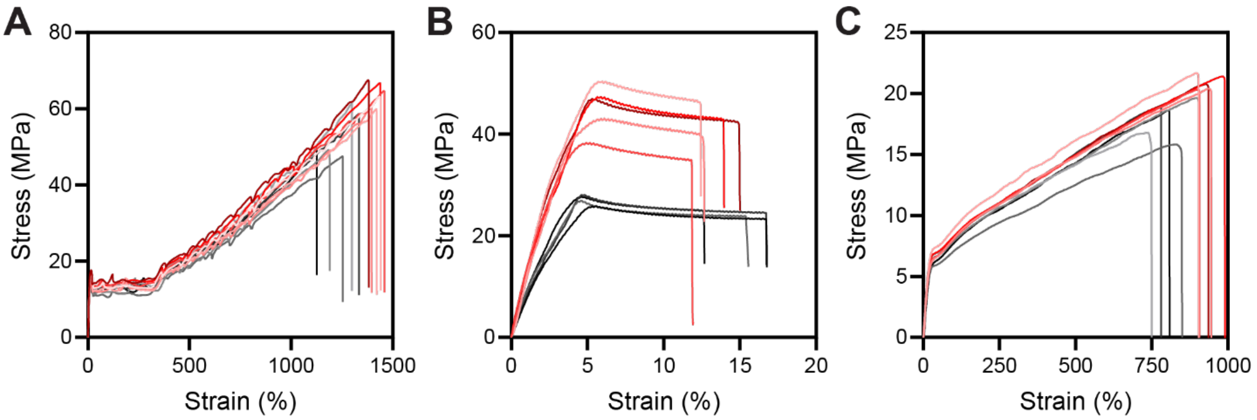

***Figure S2.*** *Stress-strain curves obtained from tensile testing of PCL (A), PLA (B) and PBAT (C) with and without spores. Neat polyesters and biocomposite polyesters are shown in grayscale and red, respectively. Toughness was calculated from the area under the curve. Ultimate tensile stress was determined by the peak stress value. Elongation at break corresponds to the strain at fracture. Young's modulus was calculated from the slope of the initial linear region.*

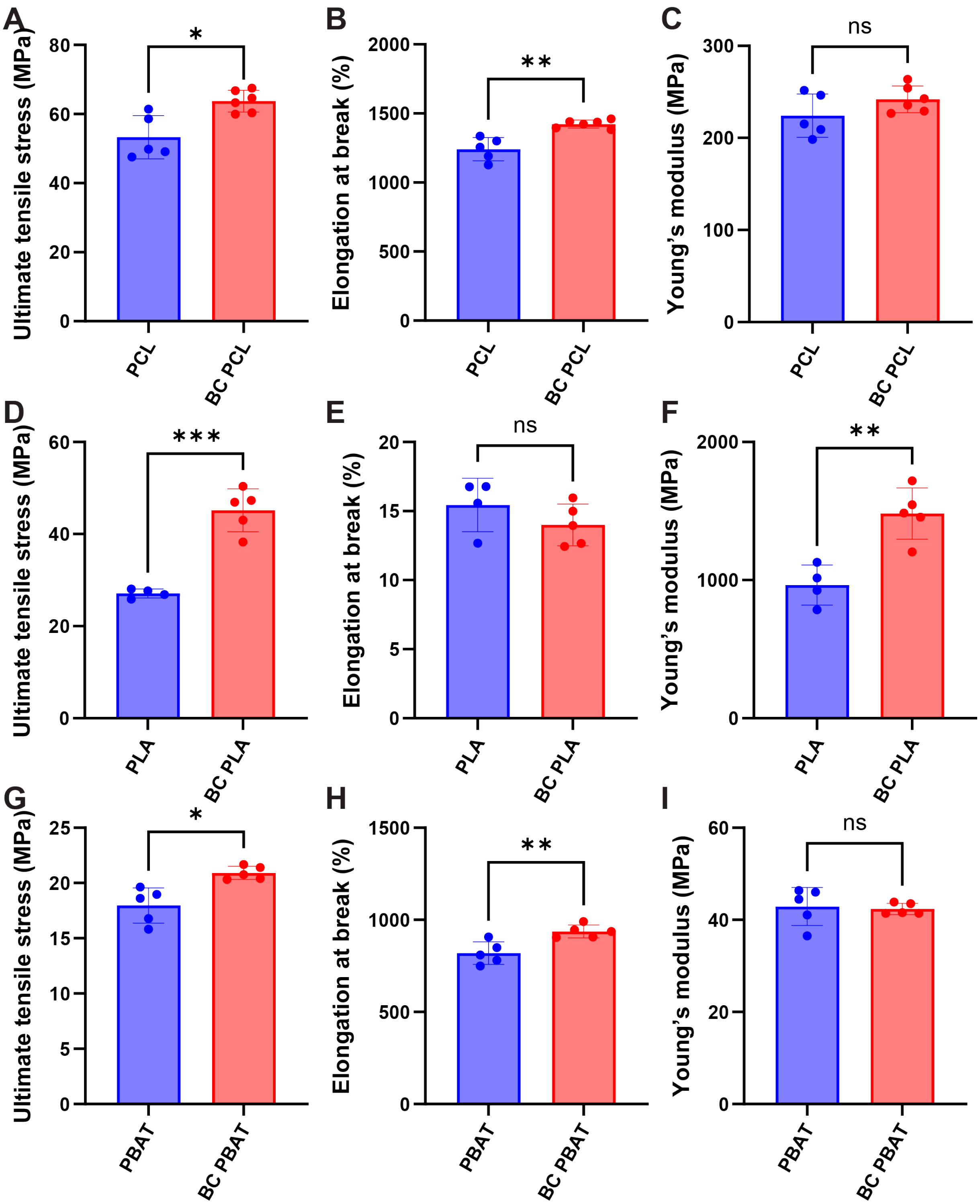

***Figure S3.*** *Tensile properties PCL (A-B), PLA (D-F) and PBAT (G-I) with and without spores. Ultimate tensile stress (A, D, G), elongation at break (B, E, H) and Young's modulus (C, F, I) are calculated from the stress vs. strain curves. Data are presented as mean values ± standard*

*deviations from four or more independent experiments. Welch's t test was performed for statistical analysis. $P < 0.05$ was considered statistically significant. Detailed indicators are as follows – ns: not significant; *$P < 0.05$; **$P < 0.01$; ***$P < 0.001$; ****$P < 0.0001$.*

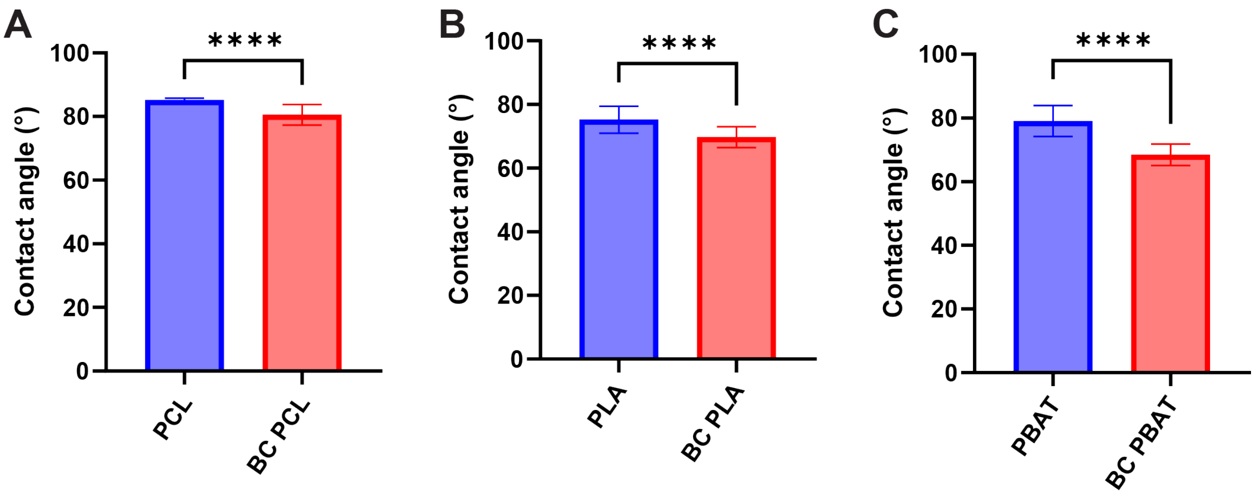

***Figure S4.*** *Water contact angle analysis of polyesters and biocomposite polyesters. (A) PCL, (B) PLA and (C) PBAT. Data are presented as mean values ± standard deviations from three independent experiments. Welch's t test was performed for statistical analysis. P < 0.05 was considered statistically significant. Detailed indicators are as follows – ns: not significant; *P < 0.05; **P < 0.01; ***P < 0.001; ****P < 0.0001.*

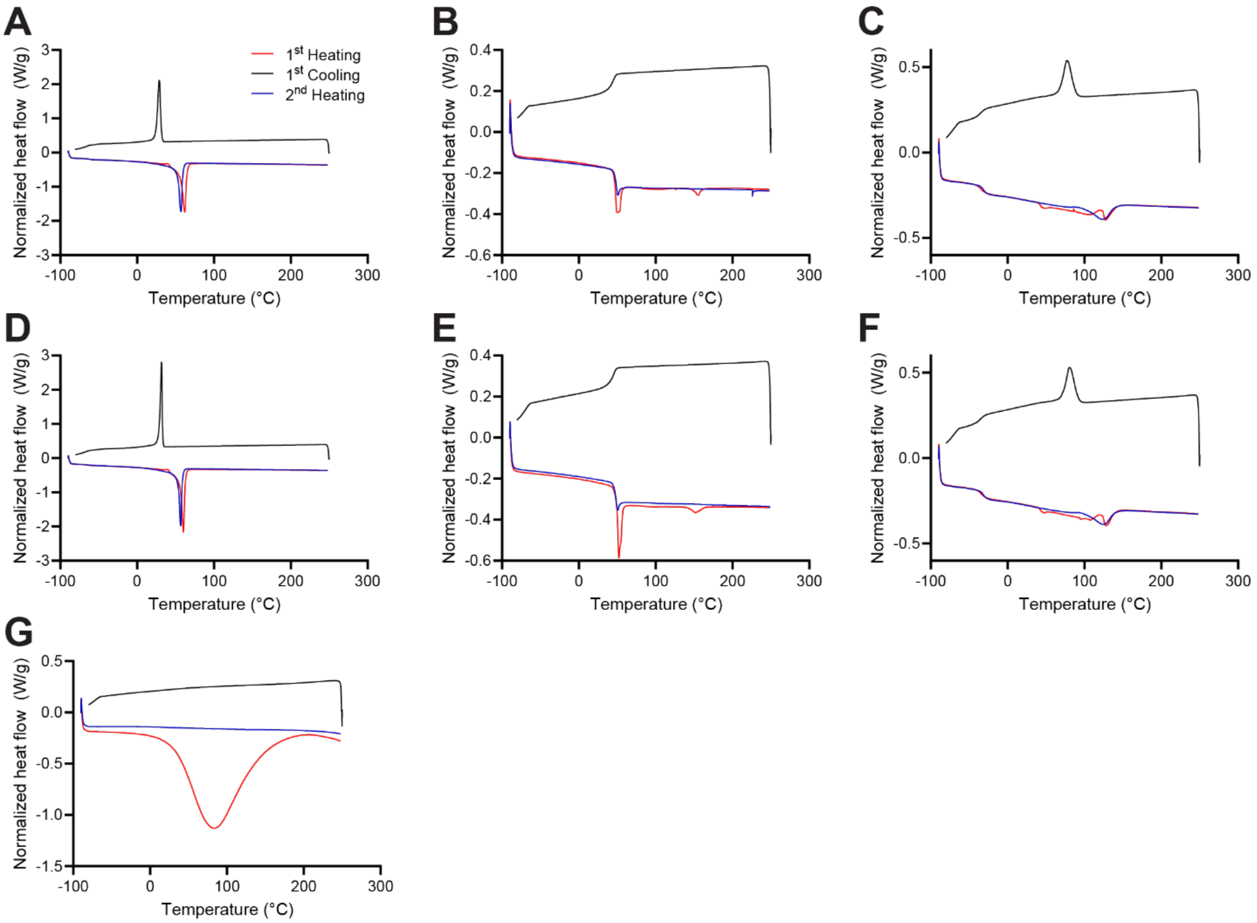

***Figure S5.*** *DSC curves of polyesters (A - C) and biocomposite polyesters (D – F). Neat and biocomposite PCL (A & D), PLA (B & E) and PBAT (C & F) showed no significant differences. DSC curves of spores are also represented (G). For all DSC analyses, red, blue and black lines indicate first heating, second heating and first cooling curves, respectively.*

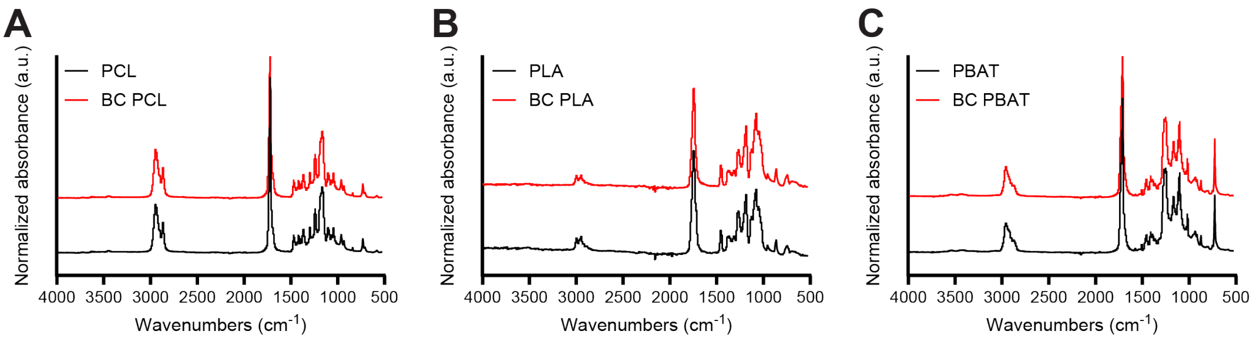

***Figure S6.*** *ATR-FTIR of polyesters and biocomposite polyesters. (A) PCL, (B) PLA and (C) PBAT.*

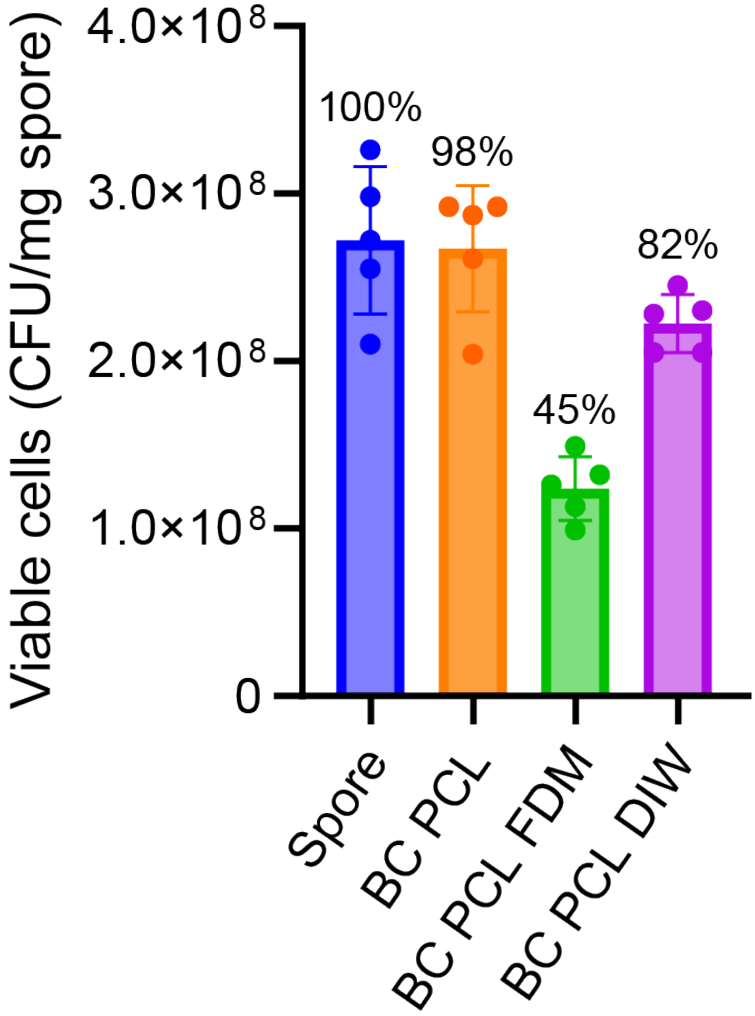

***Figure S7.*** *Viability of spores before (BC PCL) and after 3D printing via fused deposition modeling (BC PCL FDM) and direct ink writing (BC PCL DIW). Survivability was defined as relative viability, whose viability was normalized to the viability of spores treated with the same solvent extraction process (Spore). Data are presented as mean values ± standard deviations from five independent experiments.*

***Table S1.*** *Optimized extrusion conditions of polyesters. Biocomposite polyesters were prepared by adding 0.5 % (w/w) spores into the extrusion after 5 min of mixing of polyesters, followed by 15 min of mixing (5+15 min mixing time).*

| | Temperature (°C) | Mixing speed (rpm) | Mixing time (min) | Extrusion speed (rpm) |
|---|---|---|---|---|
| PCL | 65 | 36 | 5+15 | 12 |
| PLA | 135 | 36 | 5+15 | 18 |
| PBAT | 120 | 36 | 5+15 | 24 |
